# Acute Temporal Dynamics of Brain Injury Plasma Biomarkers following Controlled Football Heading

**DOI:** 10.64898/2026.09.09.749919

**Authors:** Syed Ali Muqtadir, Thomas Aston, Katy R Reid, Liivia-Mari Lember, Thomas Di Virgilio, Graeme Keith, David Hunt, Filipe Teixeira-Dias, Ferdinand Binkofski, Colin N Moran, Magdalena Ietswaart

**Affiliations:** School of Psychology, University of Stirling, Stirling, FK94LA, UK; Centre for Research and Innovation in Sport, University of Stirling, Stirling, FK94LA, UK; Institute for Infrastructure and Environment (IIE), School of Engineering, The University of Edinburgh, Edinburgh, EH93FG, UK; Institute for Neuroscience and Cardiovascular Research, University of Edinburgh, Edinburgh, EH164TJ, UK; UK Dementia Research Institute at Edinburgh University, Edinburgh, EH164SB, UK; Physiology Exercise and Nutrition Research Group, Faculty of Health Sciences and Sport, University of Stirling, Stirling, FK94LA, UK; School of Psychology and Neuroscience, University of Glasgow, Glasgow, G514TF, United Kingdom; Division of Clinical Cognitive Sciences, Medical Faculty of the RWTH Aachen University, Aachen, 52074, Germany

**Keywords:** traumatic brain injury, repeated subconcussive head impact, concussion, sport, biofluid markers, soccer

## Abstract

Footballers are routinely exposed to repeated non-concussive head impacts with (soccer) heading, the acute neurobiological consequences of which remain poorly characterised. Current evidence for acute changes in biofluid markers following heading is tempered by design limitations including inadequate impact exposure monitoring, uncontrolled confounders (e.g. exercise), the lack of a control group and heterogeneity in sampling times. This study attempts to address these limitations with a novel combination of serial sampling measuring biomarkers across five timepoints (pre and 0.5, 2, 4 and 24 h post heading), a within-subject, time-matched resting control condition and participant-level finite element modelling used for impact quantification. In a counter-balanced crossover design, twelve male football players aged 18-31 completed a heading session with 10 verified rotational headers delivered from a ball launcher at 10 m and a control condition. Impact kinematics were captured with instrumented mouthguards and modelled using the Edinburgh finite element head model (EdiFEHM). Additional monitoring measures included near point convergence (NPC) at each timepoint and the sport concussion assessment tool (SCAT6) at the pre, 0.5 and 24 h timepoints. The Quanterix Simoa N4PD Advantage Plus assay was used to quantify plasma concentration of neurofilament light (NfL), glial fibrillary acidic protein (GFAP), ubiquitin carboxy-terminal hydrolase L1 (UCH-L1) and brain-derived tau (BD-tau) measured for the first time in this context. GFAP and BD-tau showed significant main effects of time (both *P <* 0.005) but no effect of heading. UCH-L1 was excluded and NfL restricted to descriptive analysis as both fell near or below the minimum assay quantification level. NPC alongside SCAT6 symptom and cognitive scores remained unaffected by heading. Tissue strain modelling showed a mean peak MPS95 of 0.099 (range: 0.074-0.144) consistent with low magnitude non-concussive impact exposure across participants. These findings provide a methodological framework and anchor point for non-concussive impact research employing biofluid sampling while highlighting sensitivity limitations of current multiplex Simoa assays in certain biomarkers for healthy young-adult populations. Design elements including time-matched controls, serial sampling and individualised mechanical exposure outcomes are essential for characterising exposure and avoiding spurious findings. The study was preregistered with the ISRCTN registry (ISRCTN44241334).

## Introduction

Two aspects of football make it salient for the study of head impacts. The first being that it is the only sport which incorporates intentional head impacts as part of routine play and the second being the increasing popularity of the sport which is already the most widely played game in the world.^1^ The former aspect links to epidemiological studies that suggest football players are at greater risk of developing neurodegenerative conditions while the latter highlights a potentially burgeoning public health issue that needs addressing.^2–4^

It remains a challenge to define the limits of head impacts that have an influence on the brain. Head impacts resulting from football heading are classed as non-concussive, a term derived from the absence of any symptoms typically associated with concussion.^5,6^ These impacts have been associated with neurophysiological and cognitive sequelae, as well as the increased risk of neurodegenerative disease through cumulative effects.^7–10^ However, the absence of any observable markers in the acute phase makes the measurement of their effects a persistent challenge (for review see, Ntikas *et al*.^11^). Numerous methodological approaches have been adopted in extant literature spanning different cognitive domains, neuroimaging modalities and biofluid markers with heterogeneous findings.^12–16^

Blood biomarkers offer a potentially accessible approach to address this measurement challenge with technologies like single-molecule enzyme-linked immunosorbent assays (Simoa) enabling ultrasensitive protein detection.^17^ Among the most studied candidates are neurofilament light (NfL), a marker of axonal damage; glial fibrillary acidic protein (GFAP), a component of the astrocyte cytoskeleton released after astrocytic death; ubiquitin carboxy-terminal hydrolase L1 (UCH-L1), an enzyme central to neuronal repair; and tau, a protein crucial for maintaining structural integrity in axons.^18–21^ A number of studies have utilised these markers among others to investigate the effects of non-concussive head impacts but with inconsistent findings.^22–27^

A recent scoping review of 79 studies using biofluid markers to evaluate ‘repeated subconcussive impact’ exposure in sport identified several critical limitations including inadequate impact exposure monitoring, inconsistent controls for physical exertion and uncontrolled confounds such a prior head injury.^28^ The most consequential of the limitations highlighted in the review may be the heterogeneity in post-exposure sampling times and heading exposure. Most studies employ one or two post-exposure time points making it difficult to plot the response given that most blood biomarkers have distinct temporal profiles while the number of headers range from 10 to 40.^29,30^ For example within NfL literature, post-heading elevations have been reported after one hour, 24 hours, seven days and one month.^13,31–33^ On the other hand, similarly structured studies have found no change in NfL concentrations after heading.^14,22,34^ Without serial sampling across the acute post-exposure period with standardised heading exposure, there is a high risk of drawing conclusions from an incomplete picture of a biomarker’s trajectory.

The present study addresses these limitations by using a controlled heading protocol combined with serial plasma sampling across five timepoints spanning the acute post-exposure window (immediately before heading and 30 minutes, 2 hours, 4 hours and 24 hours post-heading) and a within-subject, time-matched resting control condition. Four plasma biomarkers were measured: NfL, GFAP, UCH-L1 and brain-derived tau (BD-tau), a more CNS-specific measure of total tau that has previously not been examined following controlled heading exposure.^35^ Individual heading impacts were quantified using finite element (FE) modelling, extending prior work that has characterised the biomechanical response to heading.^36,37^ This study aimed to (a) characterise the temporal profile of biomarkers’ concentrations across five timepoints following controlled heading exposure relative to a time-matched resting control, (b) examine whether FE-derived brain tissue strain was associated with individual biomarker responses in a novel concurrent characterisation of biomechanical characteristics and biological responses.

## Materials and methods

### Participants

Fourteen male football players, aged between 18 and 31 were recruited from the University of Stirling between September 2025 to November 2025. Eligibility criteria included regular active participation in football, routinely heading the ball, no history of neurological or psychiatric conditions and no diagnosed concussion in the past 12 months. A sensitivity analysis indicated that with an *n* = 12 at L = 0.05 and 80% power, the present study would be able to detect a two-tailed dependent mean difference corresponding to Cohen’s *d_z_* = 0.89. All participants provided written informed consent prior to participation. Ethical approval was obtained from the local NHS Invasive or Clinical Research Panel and all procedures aligned with guidelines set out in the Declaration of Helsinki. The study was registered with the UK Clinical Study Registry ISRCTN (ISRCTN44241334) before study commencement and the analysis plan was registered with ISRCTN before analysis commencement.

The study comprised a within-subject, counter-balanced crossover design. All participants completed three lab visits: an initial familiarisation session followed by the Heading and Control conditions in randomised order, separated by at least one week. Participants were ranked by a computer-generated random number and allocated in equal numbers to a Heading first or Control first sequence. This procedure was carried out by SAM. Familiarisation included practice with the heading apparatus, fitting of instrumented mouthguards (iMG) and measurement of anthropometric data like height and body mass. Testing for each participant was carried out at the same time of day across both conditions to control for diurnal variation observed in some biomarker concentrations.^38^

In the Heading condition, participants completed a controlled heading procedure comprising 10 rotational headers, designed to approximate routine heading practice. Balls were delivered by a Ball Launcher Trainer Pro (Net World Sports) positioned 10 m from the participant with at least 60 seconds between consecutive headers. Headers were visually verified and considered valid only if the contact exceeded (≥ 8 g) and was rated ≥ 5/10 by the participant; otherwise, the header was repeated. The full protocol can be found in Supplementary Material A. In the control condition, participants remained resting in the lab until the 4-hour sampling time with no heading exposure (Figure 1). Participants wore an iMG on the upper dentition to record head impact kinematics (Prevent Biometrics) as well as a custom skin-patch (SP) accelerometer and gyroscope (ADXL377, MPU6050) solution as a backup.

**Figure 1.**
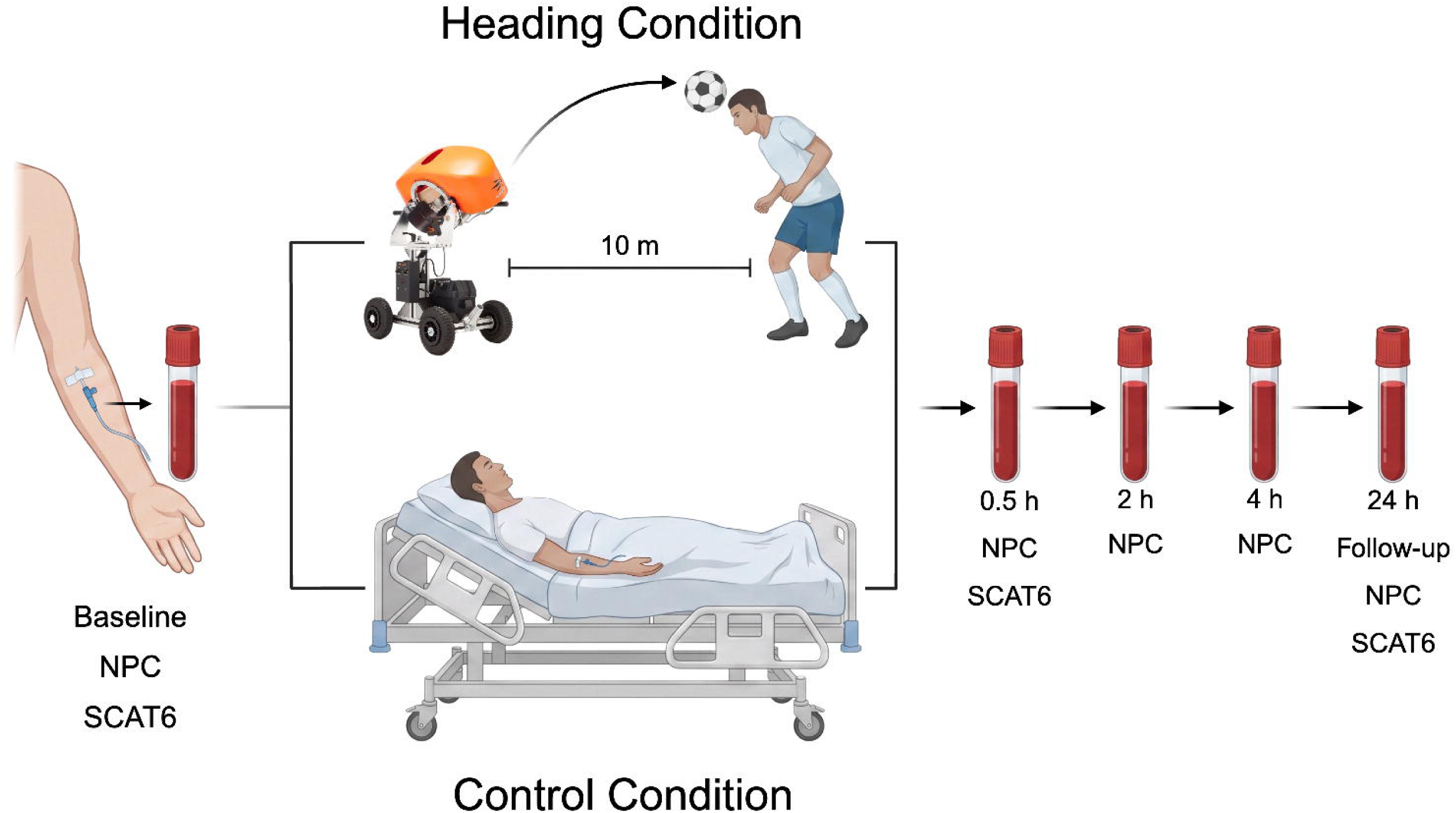
Experiment design. After baseline testing, participants stayed in a reclined resting position up to the 4 h timepoint in the control condition. The heading condition was identical to the control condition except for the addition of ten controlled heading events after baseline. A 24 h follow-up was conducted in both conditions. NPC: Near point convergence; SCAT6: Sport concussion assessment tool 6. Figure created with Biorender.com

To minimise variability across sessions, participants were asked to refrain from intense exercise as well as alcohol and recreational drugs for the 24 hours leading up to each visit. They were also provided with food and exercise diaries from their first sessions to help maintain similar food intake and activity levels across both sessions alongside instructions to hydrate with at least 500 ml of water within two hours of each session start.

### Blood sampling and laboratory analysis

Blood samples were obtained using 4 ml EDTA plasma tubes (BD vacutainer^®^) at 5 timepoints in both sessions (pre-immediate, 0.5 h, 2 h, 4 h, 24 h). Samples up to the four hour mark were obtained through an IV catheter inserted at baseline at the start of the session and standard venepuncture was used at the 24-hour follow-up. For IV catheter samples, a discard volume was collected prior to sampling and the IV line was flushed with saline after. The samples were centrifuged 6 ± 2 minutes after sampling at 1500 *g* for 15 minutes at 4 °C. Cleared plasma was aliquoted into cryovials for storage at - 80 °C for later analysis.

Sample analysis was carried out at the UK Dementia Research Institute in Edinburgh by researchers KR and SAM. Plasma biomarker concentrations were measured using a Quanterix Simoa Neurology 4-Plex D (N4PD) Advantage Plus Reagent Kits (Product No: 104822; Lot No: 504408) on a Simoa HD-X Analyzer following the manufacturer’s instructions for a 2-step immunoassay. The Simoa N4PD Advantage Plus kit measures four human brain-derived biomarkers NfL, UCH-L1, GFAP and BD-tau with manufacturer Lower Limit of Quantification and Lower Limit of Detection (LLOQs/LODs) of 1.42/0.094 pg/mL, 13.9/0.577 pg/mL, 3.22/0.121 pg/mL and 1.04/0.029 pg/mL, respectively. All samples were measured in duplicate with a 4x onboard dilution. All timepoints and conditions for a given participant were kept on the same plate. Following each run a standardised quality control (QC) pipeline was applied to the exported data. Full assay procedure and QC pipeline can be found in Supplementary Material B.

### Additional measures

The Sport Concussion Assessment Tool (SCAT6) is a standardised assessment tool used in the acute identification and evaluation of sports-related concussion.^39^ The symptom checklist and the cognitive screening from the SCAT6 were administered at the pre, 0.5 h and 24 h timepoints. In line with standard SCAT6 administration, the baseline symptom checklist was framed as how participants ‘typically felt’, while the 0.5 h and 24 h checklists were framed as how participants ‘felt now’. The cognitive screening score comprised the combined total from the orientation, immediate memory, concentration and delayed recall tasks.

Near point convergence (NPC) was measured at all timepoints. A 14-point font target on an NPC rule was held up to the tip of the participants’ nose. The target was slowly moved closer towards the participant until they reported diplopia or eye-misalignment was observed by the experimenter. The NPC measurement was repeated 3 times and averaged.

Both additional measures aimed to confirm the non-concussive nature of the heading protocol used in this study.

### Finite element modelling

The Edinburgh Finite Element Head Model (EdiFEHM) is a 50th-percentile male FEHM developed to investigate the role of intracranial surface topology, the falx and tentorium membranes and the corpus callosum on simulated brain strain response.^40^ The model has been extensively validated through the doctoral work of McGill,^40^ achieving ‘good’ biofidelity ratings as assessed by the adapted rating system of Giordano and Kleiven.^41^ For completeness, a summary of the validation results is provided in the Supplementary Material. The EdiFEHM is simulated using the LS-DYNA explicit finite element solver. It accepts linear acceleration and angular velocity along the three anatomical axes as prescribed rigid-body motion of the skull. Each simulation was run for a 100 ms duration. All simulations were performed on the Eddie compute cluster provided by the Edinburgh Compute and Data Facility (ECDF).

For each simulated impact, the EdiFEHM output was reduced to a single scalar: the volume-weighted peak 95th-percentile maximum principal strain (MPS95). To calculate this, we isolated the brain tissue elements and identified the peak strain reached by each element during the impact. These values were ranked to determine the strain level at which 95% of the total brain volume remained below. Adopting a volume-weighted 95th percentile, rather than the absolute maximum, ensures the metric reflects physical tissue deformation while mitigating the influence of numerical artifacts or anomalous single-element responses, consistent with standard practice in the literature.^42–46^ Participant averages were then calculated by averaging these peak values across all recorded headers.

### Statistical analysis

Statistical analyses were carried out using R statistical software with lme4, emmeans and lmerTest packages.^47–50^ Following QC checks (Supp. Material B), UCH-L1 was excluded entirely from the analysis as most observations (88/120) returned no quantifiable concentrations from the analyzer, a further 20 failed the CV threshold 10 returned mean concentrations below the assay LLOQ. NfL observations also saw significant attrition. Out of 120, four failed quantification from the analyzer, 37 failed the CV threshold and 10 had concentrations below the assay LLOQ. With uneven distribution of the retained observations across participants, conditions and timepoint, NfL was limited to descriptive analysis. GFAP (118/120) and BD-tau (119/120) retained sufficient data for further analysis.

The primary analysis comprised separate linear mixed models (LMM) for each biomarker (GFAP and BD-tau) with fixed effects for condition (Heading vs Control), time (pre, 0.5 h, 2 h, 4 h, 24 h) and their interaction. In line with the pre-registered analysis plan, the maximal random-effects structure that would converge was specified. For GFAP, this was a random slope for condition by participant and for BD-tau, this was a random intercept only. The primary hypothesis test for each biomarker was the condition × time interaction. Post-hoc pairwise contrasts were carried out with estimated marginal means with Holm corrections for multiple comparisons. Significance was set at an alpha of 0.05. Model residuals were inspected with Q-Q plots.

Secondary analyses comprised NPC and SCAT6 outcomes (cognitive score, symptom number, symptom severity) and were analysed with LMMs of the same structure as the primary analysis. SCAT6 symptom number and severity models were modified to account for the difference between baseline and post measures. Baseline scores averaged across the two sessions were added as a participant-level covariate.

Additional analyses were carried out to explore the link between impact kinematics and biomarker responses with the heading condition only. Averaged peak 95^th^ percentile maximum principal strain (MPS95) values for each participant, obtained from FE modelling, were used as a continuous moderator in LMMs with fixed effects of MPS95, time and their interaction, with participants as the random intercept. The MPS95 × time interaction was intended to test whether individual variation in brain tissue strain moderated biomarker temporal dynamics

## Results

Of the 14 participants recruited, two participants showed adverse reactions to venepuncture in the first instance and were withdrawn from the study. All 12 remaining participants that completed the baseline also completed every subsequent sampling timepoints, with no dropouts. Time between sessions was seven days for 11 participants, one participant’s second session was scheduled 35 days after the first one due to availability. Demographic data are presented in Table 1 and session-level data in Table 2. Individual GFAP, BD-tau and NfL levels can be found in Supplementary Material C.

**Table 1.** Sample demographics.

| Variable | Mean ± SD |
| --- | --- |
| Age (years) | 21.9 ± 3.6 |
| Height (cm) | 177.0 ± 5.5 |
| Body mass (kg) | 80.5 ± 14.3 |
| Neck circumference (cm) | 38.8 ± 5.0 |
| Head circumference (cm) | 56.8 ± 1.2 |
| Grip strength, left (kg) | 40.5 ± 5.4 |
| Grip strength, right (kg) | 41.7 ± 6.8 |
| Football experience (years) | 14.1 ± 3.6 |
| Heading experience (years) | 13.4 ± 3.7 |
| Headers per week (avg) | 12.3 ± 9.7 |
| Training hours per week (avg) | 3.7 ± 1.6 |
| Matches per week (avg) | 1.5 ± 0.7 |

**Table 2.** Session-level variables.

| Variable | Session |  |
| --- | --- | --- |
|  | Heading | Control |
| Sleep (hours) | 8.2 ± 1.0 | 8.2 ± 0.9 |
| Energy (1-10) | 7.9 ± 0.9 | 7.7 ± 1.5 |
| Stress (1-10) | 2.9 ± 2.1 | 3.1 ± 1.7 |
| Headers in the past week | 4.6 ± 4.8 | 3.8 ± 5.7 |
| Headers in the past 24 h | 0.6 ± 0.9 | 0.6 ± 0.9 |
| Ball speed (mph) | 24.8 ± 0.9 | - |
| Borg RPE | 7.7 ± 1.5 | - |

RPE: Rate of perceived exertion

### Heading exposure

All participants completed the heading protocol with 10 headers, each of which were visually verified. iMG data capture was successful for 8.4/10 ± 1.16 headers across all participants. Kinematic data are reported from iMG due to their higher accuracy relative to skin-mounted sensors.^51^ Across participants, the mean peak angular velocity (PAV) and peak linear acceleration (PLA) recorded were 11.08 ± 2.69 rad/s and 126.54 ± 19.06 m/s^2^, respectively. Mean ball speed across participants at launch was 24.8 ± 0.9 mph. Participants rated the heading activity as 7.7/20 ± 1.5 on the Borg RPE, reflecting extremely light to very light exertion. No adverse effects from the heading intervention were reported.

### Primary analysis

#### Glial fibrillary acidic protein

Two samples were excluded, one for exceeding the 20% CV threshold and the other for average enzymes/bead (AEB) being out of the calibration range. The condition × time interaction was not significant (*P* = 0.729). There was a significant effect of time, *F*(4, 85.9) = 6.38, *P <* 0.001, η^2^*_p_* = 0.23, with increased GFAP concentrations at all post-timepoints compared to baseline (all *P <* 0.018). No post-timepoints differed from each other. Mean GFAP concentrations at pre were similar across conditions (Heading: 34.9 ± 10.9 pg/ml; Control: 35.9 ± 12.4 pg/ml) and rose to comparable levels by 4 h (Heading: 41.5 ± 11.3 pg/ml; Control: 41.9 ± 10.6 pg/ml; Figure 2A and Table 3).

**Figure 2.**
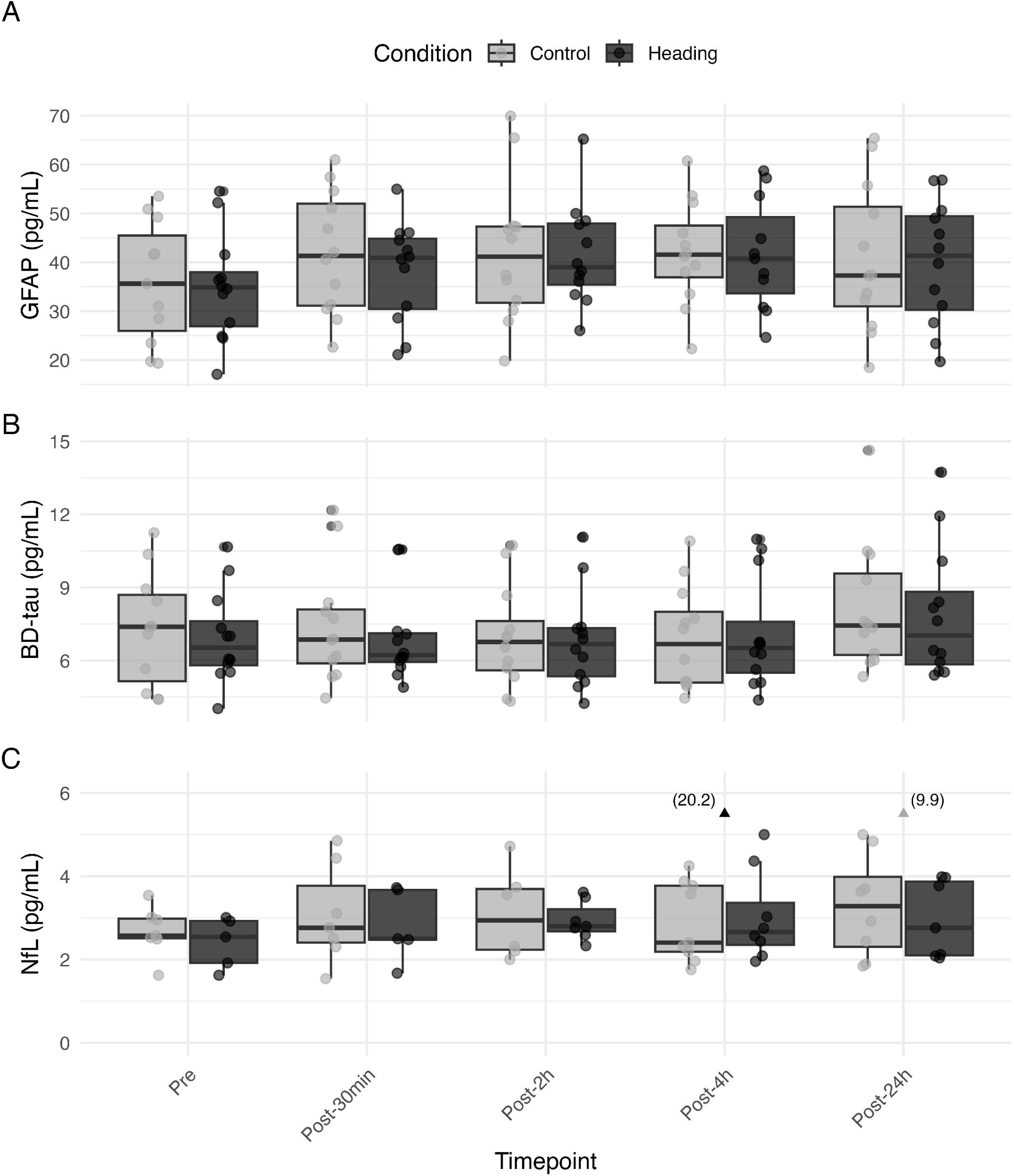
Mean biomarker concentrations for each condition across all timepoints. (**A**) GFAP concentrations (**B**) BD-tau concentrations (**C**) NfL concentrations

**Table 3.** GFAP mean concentrations.

| Condition | Time | n | Mean | SD | 95% CI Lower | 95% CI Upper |
| --- | --- | --- | --- | --- | --- | --- |
| Heading | Pre | 12 | 34.9 | 10.94 | 27.95 | 41.85 |
| Heading | 0.5h | 12 | 38.15 | 10.25 | 31.64 | 44.67 |
| Heading | 2 h | 12 | 41.56 | 10.36 | 34.98 | 48.14 |
| Heading | 4h | 11 | 41.52 | 11.29 | 33.94 | 49.1 |
| Heading | 24 h | 12 | 39.82 | 12.62 | 31.8 | 47.84 |
| Control | Pre | 11 | 35.87 | 12.45 | 27.51 | 44.24 |
| Control | 0.5 h | 12 | 41.83 | 12.5 | 33.89 | 49.78 |
| Control | 2 h | 12 | 42.14 | 14.77 | 32.76 | 51.53 |
| Control | 4 h | 12 | 41.91 | 10.56 | 35.2 | 48.62 |
| Control | 24 h | 12 | 40.81 | 15.11 | 31.21 | 50.42 |

#### Brain-Derived tau

One sample was excluded for AEB being out of the calibration range. The maximal model resulted in a singular fit so the reduced random-intercept only model was used. Residuals from the model were non-normally distributed (Shapiro-wilk, *P* = <0.001), driven by a small number of extreme upper-tail observations. Log-transformed data showed an identical pattern of results with a significant time effect (*P* = 0.005) with improved but incompletely resolved residual normality. Given the consistency of results, the untransformed model is reported here for interpretability on the original scale.

With the reduced model, the condition × time interaction was found to be non-significant (*P* = 0.884). Similar to GFAP, there was a significant effect of time, *F*(4, 98.01) = 4.18, *P* = 0.004, η^2^*_p_* = 0.15. Post-hoc comparisons showed mean concentrations at 24 h to be significantly higher than pre (*P* = 0.029), 2 h (*P* = 0.006) and 4 h (*P* = 0.012). Pre concentrations across conditions (Heading: 6.93 ± 1.89 pg/ml; Control: 7.27 ± 2.36 pg/ml) rose in both conditions by the 24 h mark (Heading: 7.92 ± 2.72; Control: 8.17 ± 2.64; Figure 2B and Table 4).

**Table 4.** BD-tau mean concentrations.

| Condition | Time | n | Mean | SD | 95% CI Lower | 95% CI Upper |
| --- | --- | --- | --- | --- | --- | --- |
| Heading | Pre | 12 | 6.93 | 1.89 | 5.73 | 8.13 |
| Heading | 0.5 h | 12 | 6.9 | 1.83 | 5.74 | 8.07 |
| Heading | 2 h | 12 | 6.83 | 1.98 | 5.57 | 8.08 |
| Heading | 4 h | 12 | 7.05 | 2.25 | 5.62 | 8.48 |
| Heading | 24 h | 12 | 7.92 | 2.72 | 6.19 | 9.65 |
| Control | Pre | 11 | 7.27 | 2.36 | 5.69 | 8.86 |
| Control | 0.5 h | 12 | 7.41 | 2.37 | 5.91 | 8.92 |
| Control | 2 h | 12 | 6.95 | 2.09 | 5.62 | 8.27 |
| Control | 4 h | 12 | 6.89 | 2.1 | 5.55 | 8.23 |
| Control | 24 h | 12 | 8.17 | 2.64 | 6.49 | 9.85 |

#### Neurofilament light

With 69 surviving observations, NfL concentrations showed no consistent temporal pattern across the two conditions from pre (Heading: 2.40 ± 0.61 pg/ml; Control: 3.68 ± 0.60 pg/ml) to 24 h (Heading: 2.96 ± 0.92 pg/ml; Control: 3.89 ± 2.62 pg/ml; Figure 2C and Supp. Table 3).

### Secondary analysis

#### Near point convergence

The condition × time interaction for NPC was not significant (*P* = 0.869). Values remained stable across both sessions between pre (Heading: 8.58 ± 2.53 cm; Control: 9.03 ± 2.20 cm), 4 h (Heading: 9.08 ± 2.42 cm; Control: 9.44 ± 2.18 cm) and 24 h (Heading: 9.44 ± 2.71 cm; 9.61 ± 2.24 cm; Figure 3A).

**Figure 3.**
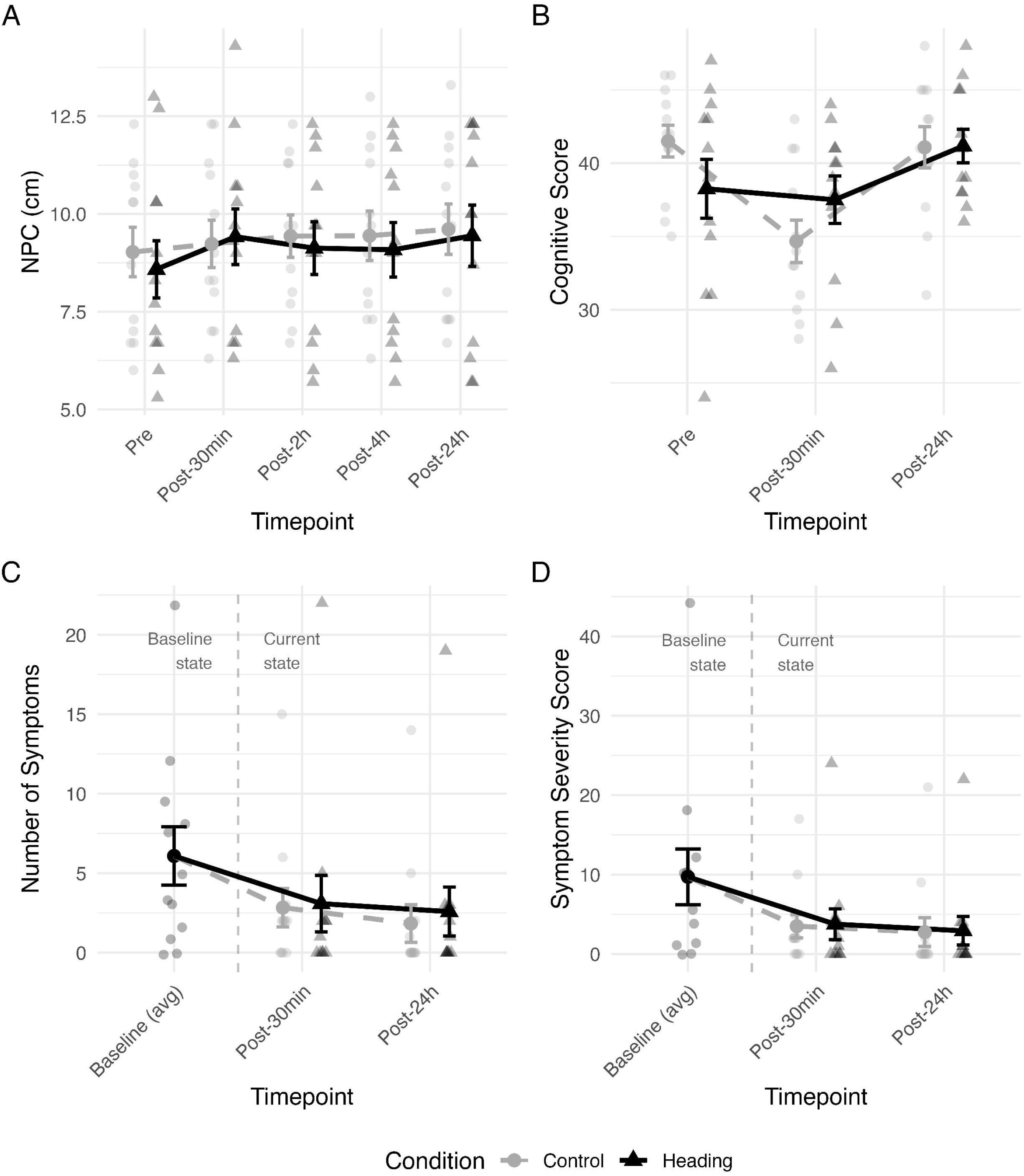
Secondary measure trends over time and across conditions. **(A)** Near point convergence **(B)** Cognitive score **(C)** SCAT6 number of symptoms **(D)** SCAT6 symptom severity scores

#### SCAT6 cognitive screening

A significant condition × time interaction was found *F*(2, 55) = 3.415, *P* = 0.04. However, post-hoc pairwise contrasts between conditions at each timepoint did not reach significance (pre: *P* = .054, 0.5 h: *P* = 0.091, 24 h: *P* = 0.960). The interaction appeared to be driven by a decrease in Control scores at 0.5 h (41.5 ± 3.75 to 34.7 ± 5.04), a change that was not reflected in the Heading condition (38.2 ± 6.96 to 37.5 ± 5.62; Figure 3B).

#### SCAT6 symptom evaluation

Neither SCAT6 symptom number nor symptom severity showed any significant condition × time interactions (symptom number: *P* = 0.550; symptom severity: *P* = 0.941; Figure 3C and 3D).

#### FE Modelling

Peak MPS95 from finite element modelling of all headers for each participant averaged 0.099, ranging from 0.074 to 0.144 (Figure 4). MPS95 x time interactions were not significant for either GFAP (*P* = 0.389) or BD-tau (*P* = 0.787).

**Figure 4.**
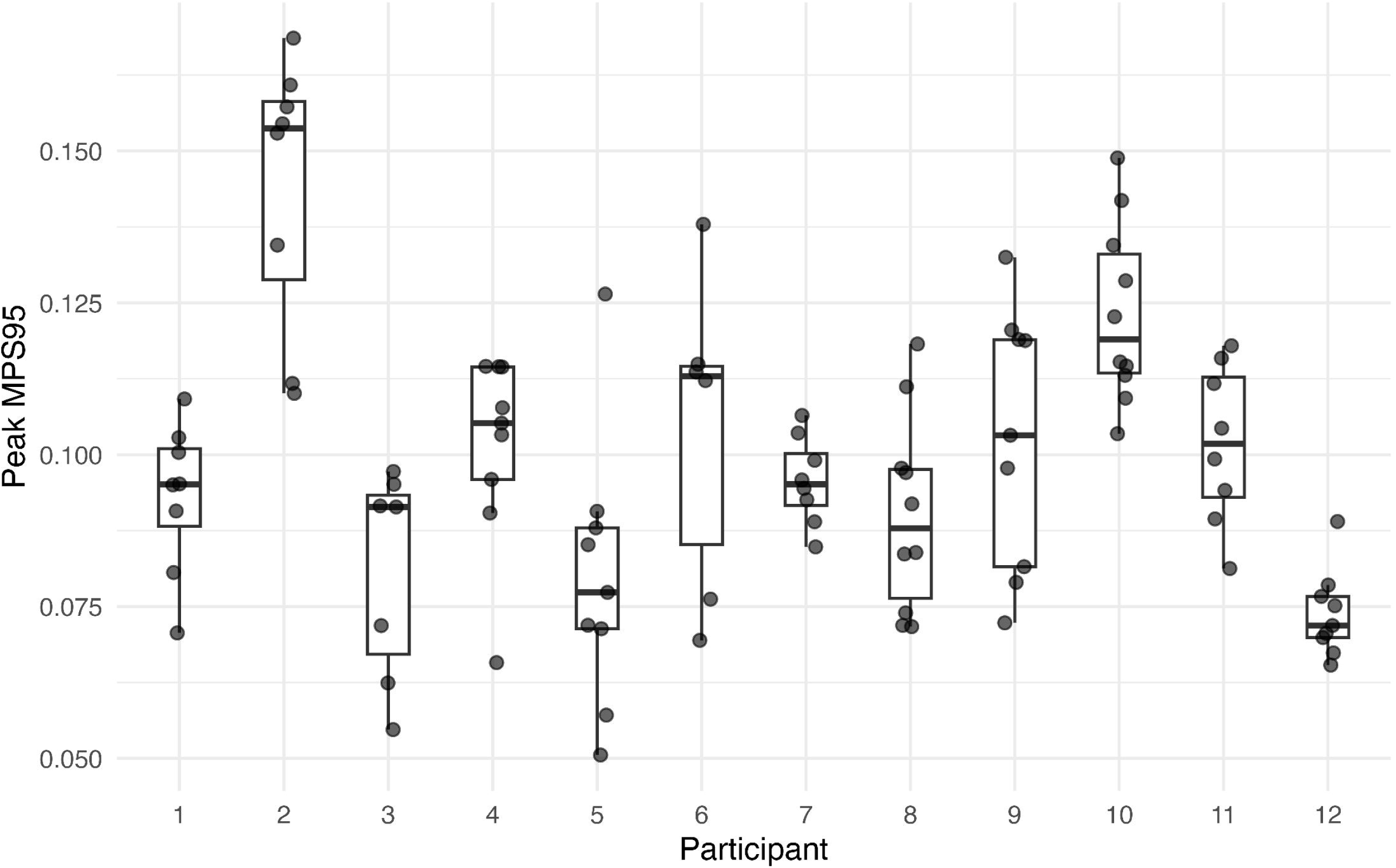
Individual MPS95 values. All instrumented mouthguard-recorded impacts across participants.

## Discussion

This study examined the acute effects of repeated non-concussive head impacts on plasma biomarkers of neuronal and glial injury using serial sampling across five timepoints in a within-subjects crossover design. Primary analysis showed no significant changes in GFAP or BD-tau plasma concentrations following 10 rotational headers relative to a time-matched resting control condition. NfL and UCH-L1 could not be meaningfully evaluated due to substantial data loss at the quality control stage. Monitoring outcomes, namely, NPC and SCAT6 symptom measures showed no significant heading effects with the exception of a nominally significant condition × time interaction observed for SCAT6 cognitive scores was driven by the control condition and did not survive corrections for multiple comparisons. Exploratory finite element modelling revealed no relationships between individual brain tissue strain magnitude and biomarker (GFAP and BD-tau) responses. Nonetheless, these findings should not be read as a reassurance about the safety of heading. As we discuss below, the pattern of results highlights the importance of methodological considerations for biomarker investigations including time-matched control conditions and the analytical limits of current multiplex Simoa assays in healthy young-adult populations.

Although GFAP concentrations rose significantly across timepoints, no difference was observed between Heading and Control conditions. The time-matched control condition was therefore critical to the interpretation of these findings as without it, the significant main effect of time could plausibly have been misattributed to a heading-related response rather than to temporal processes unrelated to the intervention.^38^ The null condition × time interaction for GFAP is consistent with prior interventional heading studies which have reported no acute GFAP changes in either CSF or plasma.^14,52^ GFAP elevations previously reported following non-concussive impacts are associated with higher mechanical loads than those in the present study. GFAP elevations have been reported in American football players following head impacts with PLA values of 27.7 ± 17 g, substantially higher than the 12.9 ± 1.94 g recorded here.^53^ Notably, GFAP elevations and time courses may depend on injury severity. Mild and transient elevations have been found in serum GFAP up to 48 hours following non-concussive head trauma but marked increases with concussion where it was detectable within one hour and remained elevated for seven days.^54^ GFAP may have limited utility in detecting acute effects at exposures of the magnitude present in the current study though this does not preclude sensitivity at higher impact or cumulative mechanical loads.

To our knowledge, this is the first study to track BD-tau responses to controlled football heading. BD-tau is a selective measure of CNS derived tau with greater specificity than total tau as a measure of neuronal injury.^35^ BD-tau concentrations have been shown to scale with injury severity and may serve as a more reliable marker for traumatic brain injury (TBI) recovery.^55^ The null interaction for BD-tau indicates that even a CNS-specific measure of tau was not able to capture any heading related changes at the current exposure levels. Studies employing peripheral tau found its levels unchanged following 40 headers or 10 headers.^13,52^ It should be noted that peripheral t-tau shows poor association with CSF t-tau and may be susceptible to exercise effects.^22,56,57^ While the current findings cannot be directly compared to existing t-tau findings in the football heading literature, it appears that despite its CNS specificity, BD-tau did not show any measurable changes after 10 controlled headers.

For two of our four target analytes, UCH-L1 and NfL, inferential analysis was precluded by data loss at the quality control stage. In both cases, the pattern of attrition points to assay sensitivity, rather than experimental confounds, as the primary cause. Although the specific limitations differ between the two markers. In the case of UCH-L1, the 4-Plex D UCH-L1 LLOQ of 13.9 pg/ml is substantially higher than that of the other analytes in the same kit (GFAP, 3.22 pg/ml; NfL, 1.42 pg/ml; BD-tau, 1.04 pg/ml). The even distribution of NaN and below-LLOQ values across timepoints and conditions indicated that the data loss was driven by assay performance rather than experimental manipulation. This issue has been reported in other studies, where 47.2 % of baseline UCH-L1 concentrations fell below the Simoa 4-plex B assay LLOQ of 9.38 pg/ml in a sample of collegiate athletes.^58^ In addition, UCH-L1 concentrations are further complicated by endogenous factors including sex and exercise effects, with male adolescents showing lower baseline levels than females and a negative correlation between peripheral UCH-L1 and testosterone.^59,60^ The combination of the elevated assay quantification floor and confounding influences makes UCH-L1 an unreliable candidate for non-concussive impact research using current multiplex platforms.

Similar to UCH-L1, data loss in NfL appears to also be driven by the proximity of healthy young-adult NfL concentrations to the assay’s sensitivity floor. The mean NfL concentration in surviving observations across all timepoints and conditions was 3.23 ± 2.39 pg/ml which is nearly a full standard deviation below the mean reported for healthy adults aged 18 to <51 (6.07 ± 2.85 pg/ml) and is closer to the mean reported for the 5 to <18 age group (3.89 ± 1.67 pg/ml).^61^ This is consistent with findings from the United States where a median of 4.2 pg/ml (95% CI: 3.9-4.4) for healthy adults aged 20-29 was reporting using a singleplex Simoa NF-Light Advantage assay.^62^ In the context of heading, a singleplex Simoa NF-Light HD-1 assay (LLOQ: 0.58 pg/ml) was used to detect significant NfL elevations at one hour and one month after exposure to 40 headers.^13^ Of note here is the use of singleplex NfL assays. No published study has directly compared singleplex and multiplex Simoa performance at the lower end of the dynamic range where a healthy young population’s NfL concentrations would sit. The multiplex-singleplex comparison that does exist was conducted with acute TBI patients with reported NfL concentrations that were well within the dynamic range of both assays.^63^ Whether the 4-Plex D retains that analytical equivalence with single-plex NfL assays at lower concentrations of NfL remains an open question. Future studies with healthy young cohorts may use assay optimisation to improve quantification. Reducing the sample dilution from the manufacturer recommended 4x may bring NfL AEBs onto the standard curve, however, this must be balanced against an increased susceptibility to non-specific binding.

Secondary outcomes were consistent with biomarker findings. NPC did not differ between conditions, a finding that fits with a broader pattern of mixed results in the literature. Studies employing similar 10 header protocols have produced mixed outcomes. Mixed findings are present in extant literature with acutely worsening NPC up to 24 hours reported,^64,65^ while no such effects have been found in studies with adolescents.^66,67^ As with blood biomarkers, the heterogeneity in the literature with different impact kinematics and experimental designs makes it difficult to identify a causal relationship between repeated non-concussive head impacts and NPC. SCAT6 outcomes showed no clinically meaningful heading related changes confirming that the participants were exposed to a controlled mechanical stimulus without the clinical sequelae associated with concussion.

The mechanical load experienced by participants in the current sample was indexed using FE-derived MPS95, an estimate of brain deformation which captures tissue-level effects that kinematics variables like peak linear or rotational acceleration may not. In American Football literature, MPS95 has been associated with longitudinal changes in UCH-L1 and related brain strain metrics have outperformed kinematic metrics in accounting for pre- and post-season DTI changes.^25,68^ While strain-based injury markers remain model specific, several attempts have been made to identify MPS95 thresholds for the risk of brain injury (mild TBI) ranging from 0.27 to 0.23.^69,70^ The strain magnitudes in the current sample (0.099 ± 0.019) were well below concussion thresholds reported in extant literature. The values were comparable to those reported from a laboratory protocol with varying ball speeds in adult amateur players and a different FE model (0.096 ± 0.252) but exceeded those recorded from adolescent players (0.045-0.048).^36,67^ No moderating effects of MPS95 on GFAP or BD-tau trajectories were observed. The narrow exposure range in the present sample, while demonstrating the consistency of impacts derived from the controlled heading protocol, limits the interpretation of current findings to a bound range of heading exposure.

This study, while constrained to a small sample size of 12 and limited to the detection of large effects, nuances utility issues that are critical considerations for future studies. Given the nature of the decisively null results for GFAP and BD-tau, a larger sample size is unlikely to yield contradictory findings. Future studies applying the methodological framework presented here may critical insights with other biomarkers within the sensitivity-related constraints documented here. Sex differences have been documented for several of the target analytes, making an extension to female players a priority for future work, a male-only sample in the present work limits generalisability.^59,71^ Our within-subjects crossover design helps mitigate but not eliminate the limitations arising from a small sample size by accounting for any between-person variance in baseline biomarker concentrations and other physiological factors. Data on normal within-person variance in biofluid marker levels in healthy populations remains limited, which warrants further study. The laboratory heading protocol, while lacking the unpredictability of match conditions, allowed us to control the impact characteristics which was essential for linking biomarker responses to a mechanical stimulus while controlling for extraneous factors like exercise effects. A recent study reporting acute elevations in p-tau-217 and S100B after match heading exposure illustrates the difficulty of isolating heading effects in naturalistic settings.^16^ Their adjustment for exercise, while methodologically careful, relied on a single composite that captured just over half of the variance in exercise measures making confounding by exercise related factors difficult to fully exclude. The present study sidesteps the former problem by eliminating exertion and contact entirely through a within-subject resting control condition, allowing observed differences to be attributed to the heading exposure itself. Additionally, the first and only systematic review on the topic showed S100B to be elevated by exercise alone and thus unsuited to indexing heading impact effects, underscoring the importance of biomarker selection.^28^ While the loss of NfL and UCH-L1 data restricts the current study to GFAP and BD-tau inferences, the nature of the data loss and its link to assay performance in healthy young adults highlights a gap within extant literature that merits further exploration.

Determining whether repeated heading induces acute neurobiological change remains an open question and the present findings should not be interpreted as evidence that it does not. The absence of heading-specific effects on GFAP and BD-tau time courses may reflect a sub-threshold mechanical stress while any effects on NfL and UCH-L1 are masked by assay sensitivity limitations. The present study contributes to a methodological framework for approaching investigations into non-concussive impacts with biomarkers highlighting the importance of time-matched control condition and serial sampling. Future works would be well served by combining more sensitive assay platforms, impact modelling and serial sampling to parse out any genuine impact-specific effects from natural biological variation. Until such data are available the question of acute neural injury following football heading remains in doubt.

## Data availability

The authors confirm that the data supporting the findings of this study are available within the article and its supplementary material.

## Funding

This work was supported through funding by Medical Research Scotland and a RS MacDonald Facility Access grant administered by SINAPSE and SULSA awarded to SAM, TA and GK.

## Competing interests

The authors report no competing interests.

## Notes

### Competing Interest Statement

The authors have declared no competing interest.

