## Supplementary Material A; Supplementary Material B; Supp. Material B; Supplementary Material C; Supp. Table 3 for "Acute Temporal Dynamics of Brain Injury Plasma Biomarkers following Controlled Football Heading"

##### Protocol for Controlled Heading Activity

Adapted from Bevilacqua *et al.*[1] and Di Virgilio *et al.*[2], this protocol pertains to standing rotational headers.

1. **Setup**
    a. The ball machine is placed 10 meters away from the participant.
    b. The standard size 5 football (68-70 cm circumference) is inspected for surface defects and inflated to the required pressure level (10 psi).
    c. The participant is briefed on the task. It is ensured that the participant understands the following.
        i. The ball machine will project the football to them.
        ii. They are required to perform standing linear headers i.e. making contact with the ball with only their forehead and directing it back towards where the ball is projected from. They are to avoid jumping or moving excessively for the duration of the task.
        iii. The activity will begin with the participants catching the ball with their hands in front of their forehead.
    d. Kinematic measurement devices are affixed to the participant (if applicable)
2. **Familiarisation**
    a. The participant is positioned 10 metres from the ball machine.
    b. The ball machine is turned on and both dials on the control panel are set to the desired speed.
    c. The participant verbally confirms that they are ready to catch the ball with their hands
    d. The football is loaded into the ball machine and launched.

e. The participant is asked to confirm if they find the speed of the ball to be acceptable.

f. The launch speed and launch angle are adjusted if needed.

g. Steps 3 to 6 are repeated until the participant feels comfortable to start the heading activity.

3. **Heading**

    a. Ensure kinematic measurement devices are running.

    b. The ball machine is turned on and both dials on the control panel are set to the desired speed.

    c. The participant verbally confirms that they are ready to head the ball.

    d. The football is loaded into the ball machine and launched.

    e. After each header, the accelerometer is checked to ensure that the contact was recorded.

    f. The participant is asked to rate each header contact out of 10. Any header that glances off the participants' head, is missed, or is rated by the participant as less than 5 is repeated.

    g. Steps 3 and 6 are repeated with 1 minute in between each header.

    h. After 10 heading impacts have been recorded, the ball machine is turned off. The accelerometer is also turned off and removed.

    i. Participants are asked to rate the intensity of the heading activity on the Borg Rating of Perceived Exertion (RPE)

#### Supplementary Material B

##### Quanterix Simoa N4PD Advantage Plus assay

Plasma samples were allowed to rise to room temperature for one hour after which they were briefly vortexed and centrifuged at 10,000 *g* for 5 minutes to remove debris and to separate out the lipid layer. Plasma (95 µl) was then transferred onto Quanterix 96-well plates in pseudo-randomised order wherein all conditions and timepoints for a participant were kept on the same plate (within the same assay run). The running order was set after two-person verification of each sample. After the addition of calibrators and controls, the plates were then sealed with Pierce XP-100 plate films and loaded into the HD-X analyzer. All samples were measured in duplicate with a 4x on-board dilution. Calibrators were measured neat in triplicate. Digital, analogue and in-house controls were run in duplicate with a 4x on-board dilution. In step 1, the 100 µl of each diluted sample (25 µl plasma and 75 µl N4PD plasma sample diluent) was incubated with 25 µl of paramagnetic beads coated with target antibodies and 20 µl of a biotinylated detector for 35:15 minutes. This was followed by the addition of 100 µl of N4PD Advantage Plus streptavidin-beta-galactosidase (SBG) and further incubation lasting 5:15 minutes in step 2. Finally, for measurement, 50 µl of resorufin beta-D-galactopyranoside (RGP) was added and the solution transferred to an array on a Simoa disc. The disc was then sealed with Simoa HD-X sealing oil and an image taken after 30 seconds to measure the fluorescent product from the hydrolysis of RGP by SBG-bound captured targets.

##### Quality control checks

After each run, quality control checks were carried out, starting with the batch calibration report generated by the HD-X Analyzer. Here, the presence of all replicates for each standard was confirmed and the line of best fit was evaluated alongside checks for any flags or errors. This was followed by data quality checks run on the exported assay data comprising the following steps in order: bead number consistency check, checking kit Control 1 (digital) and kit Control 2 (analogue) mean concentrations were within the expected range for the kit and lot-specific Certificate of Analysis (CoA), checking all CVs were below 20% for each sample and finally checking that sample mean concentrations were above the Lower Limit of Quantification (LLOQ) and Lower Limit of Detection (LOD) for each biomarker.

### Supplementary Material C

The following tables contain all biomarker concentration data used in this study.

**Supplementary Table 1 Individual GFAP concentration means**

| ID | Condition | Pre | 0.5 h | 2 h | 4 h | 24 h |
|---|---|---|---|---|---|---|
| 1 | Heading | 27.62 | 28.61 | 36.1 | 40.75 | 31.15 |
| 2 | Heading | 24.49 | 22.53 | 37.41 | 36.51 | 23.36 |
| 3 | Heading | 41.56 | 46.05 | 49.99 | <NA> | 42.9 |
| 4 | Heading | 35.17 | 38.86 | 44 | 44.86 | 49.02 |
| 5 | Heading | 52.21 | 54.98 | 65.23 | 57.25 | 56.72 |
| 6 | Heading | 33.58 | 42.47 | 48.5 | 53.67 | 56.85 |
| 7 | Heading | 17.08 | 21.1 | 26.03 | 24.63 | 19.67 |
| 8 | Heading | 36.38 | 41.14 | 33.4 | 37.7 | 39.77 |
| 9 | Heading | 54.54 | 44.51 | 38.24 | 58.73 | 50.61 |
| 10 | Heading | 24.82 | 31.06 | 32.26 | 30.77 | 27.63 |
| 11 | Heading | 34.59 | 45.9 | 47.77 | 41.76 | 45.79 |
| 12 | Heading | 36.76 | 40.63 | 39.75 | 30.11 | 34.4 |
| Mean [95% CI] | | 34.9 [27.95, 41.85] | 38.15 [31.64, 44.67] | 41.56 [34.98, 48.14] | 41.52 [33.94, 49.1] | 39.82 [31.8, 47.84] |
| 1 | Control | 23.48 | 30.49 | 27.94 | 38.05 | 26.93 |
| 2 | Control | 28.48 | 35.53 | 36.37 | 41.27 | 32.35 |
| 3 | Control | 50.89 | 57.44 | 65.5 | 60.77 | 65.42 |
| 4 | Control | 41.74 | 51.13 | 37.38 | 43.45 | 37.24 |
| 5 | Control | 53.51 | 61 | 69.95 | 53.62 | 63.72 |
| 6 | Control | 41.59 | 46.87 | 47.27 | 45.96 | 43.31 |
| 7 | Control | 19.71 | 22.63 | 19.8 | 22.3 | 18.48 |
| 8 | Control | 19.34 | 28.31 | 32.21 | 39.34 | 49.95 |
| 9 | Control | 49.25 | 54.62 | 47.42 | 52.27 | 55.72 |
| 10 | Control | 31 | 31.32 | 46.79 | 33.51 | 25.62 |
| 11 | Control | 35.63 | 42.09 | 44.94 | 41.86 | 33.66 |
| 12 | Control | <NA> | 40.57 | 30.18 | 30.47 | 37.37 |
| Mean [95% CI] | | 35.87 [27.51, 44.24] | 41.83 [33.89, 49.78] | 42.14 [32.76, 51.53] | 41.91 [35.2, 48.62] | 40.81 [31.21, 50.42] |

**Supplementary Table 2 Individual BD-tau concentration means**

| ID | Condition | Pre | 0.5 h | 2 h | 4 h | 24 h |
|---|---|---|---|---|---|---|
| 1 | Heading | 6.06 | 6.14 | 6.15 | 6.68 | 8.17 |
| 2 | Heading | 5.48 | 7.09 | 7.12 | 6.27 | 5.94 |
| 3 | Heading | 5.53 | 5.41 | 4.93 | 5.64 | 5.4 |
| 4 | Heading | 9.7 | 10.55 | 11.07 | 10.12 | 10.08 |
| 5 | Heading | 7.01 | 6.14 | 6.89 | 6.73 | 7.63 |
| 6 | Heading | 10.67 | 10.57 | 9.81 | 10.59 | 13.73 |
| 7 | Heading | 4.03 | 4.9 | 5.13 | 4.38 | 5.55 |
| 8 | Heading | 7 | 5.75 | 4.24 | 5.05 | 11.93 |
| 9 | Heading | 8.46 | 6.82 | 7.31 | 10.98 | 8.4 |
| 10 | Heading | 6.04 | 6.3 | 6.46 | 6.35 | 6.42 |
| 11 | Heading | 5.89 | 6 | 5.43 | 5.1 | 6.29 |
| 12 | Heading | 7.33 | 7.2 | 7.38 | 6.75 | 5.52 |
| Mean [95% CI] | | 6.93 [5.73, 8.13] | 6.9 [5.74, 8.07] | 6.83 [5.57, 8.08] | 7.05 [5.62, 8.48] | 7.92 [6.19, 9.65] |
| 1 | Control | 8.45 | 6.18 | 6.56 | 7.75 | 7.14 |
| 2 | Control | 7.39 | 7.74 | 7.28 | 7.55 | 7.6 |
| 3 | Control | 5.67 | 6.04 | 5.98 | 4.94 | 7.36 |
| 4 | Control | 11.25 | 12.18 | 10.41 | 10.91 | 10.49 |
| 5 | Control | 7.09 | 6.88 | 7 | 6.05 | 7.51 |
| 6 | Control | 10.36 | 11.52 | 10.73 | 9.66 | 14.63 |
| 7 | Control | 4.64 | 5.42 | 4.42 | 4.46 | 5.34 |
| 8 | Control | 4.42 | 4.46 | 4.33 | 5.11 | 10.37 |
| 9 | Control | 8.94 | 8.37 | 6.98 | 8.76 | 9.3 |
| 10 | Control | 7.4 | 8.01 | 8.67 | 7.3 | 6.3 |
| 11 | Control | 4.4 | 5.34 | 5.35 | 5.04 | 5.94 |
| 12 | Control | <NA> | 6.84 | 5.68 | 5.14 | 6.02 |
| Mean [95% CI] | | 7.27 [5.69, 8.86] | 7.41 [5.91, 8.92] | 6.95 [5.62, 8.27] | 6.89 [5.55, 8.23] | 8.17 [6.49, 9.85] |

**Supplementary Table 3 Individual NfL concentration means**

| ID | Condition | Pre | 0.5 h | 2 h | 4 h | 24 h |
|---|---|---|---|---|---|---|
| 1 | Control | <NA> | <NA> | 3.74 | 3.88 | 3.7 |
| 2 | Control | 3.02 | 2.76 | <NA> | 3.58 | 3.64 |
| 3 | Control | <NA> | <NA> | <NA> | <NA> | <NA> |
| 4 | Control | 2.95 | 3.11 | 2 | 2.32 | 2.44 |
| 5 | Control | 1.62 | 1.54 | 2.32 | <NA> | 1.84 |
| 6 | Control | 3.54 | 4.43 | 4.72 | 4.25 | 4.84 |
| 7 | Control | 2.49 | 4.85 | <NA> | 1.76 | <NA> |
| 8 | Control | <NA> | <NA> | <NA> | 1.96 | 1.89 |
| 9 | Control | <NA> | <NA> | <NA> | 3.78 | 9.88 |
| 10 | Control | 2.53 | 2.31 | 3.56 | 2.41 | <NA> |
| 11 | Control | 2.57 | 2.51 | 2.2 | 2.19 | 2.92 |
| 12 | Control | <NA> | <NA> | <NA> | <NA> | <NA> |
| Mean<br>[95% CI] | | 2.68<br>[2.13, 3.23] | 3.07<br>[1.98, 4.17] | 3.09<br>[1.95, 4.23] | 2.9<br>[2.17, 3.63] | 3.89<br>[1.7, 6.09] |
| 1 | Heading | 2.92 | 2.5 | 3.5 | 3.03 | 3.99 |
| 2 | Heading | 1.92 | 3.73 | 2.92 | 2.57 | 2.09 |
| 3 | Heading | <NA> | <NA> | <NA> | <NA> | <NA> |
| 4 | Heading | 1.62 | 2.48 | 2.59 | 2.75 | 2.76 |
| 5 | Heading | <NA> | <NA> | 2.33 | 1.96 | <NA> |
| 6 | Heading | 3.01 | 3.67 | 3.62 | 4.36 | 3.77 |
| 7 | Heading | <NA> | 1.67 | <NA> | <NA> | 2.12 |
| 8 | Heading | <NA> | <NA> | <NA> | <NA> | <NA> |
| 9 | Heading | <NA> | <NA> | <NA> | 20.2 | <NA> |
| 10 | Heading | 2.54 | <NA> | 2.76 | 2.44 | 2.04 |
| 11 | Heading | <NA> | <NA> | 2.8 | 2.09 | 3.97 |
| 12 | Heading | <NA> | <NA> | <NA> | <NA> | <NA> |
| Mean<br>[95% CI] | | 2.4<br>[1.64, 3.16] | 2.81<br>[1.72, 3.9] | 2.93<br>[2.5, 3.36] | 4.92<br>[-0.27, 10.12] | 2.96<br>[2.11, 3.81] |

#### Supplementary Material D

This appendix summarises the validation of the Edinburgh Finite Element Head Model (EdiFEHM), as reported in the doctoral thesis of McGill.[3] The EdiFEHM used corresponds to the 'All' model variant described in that work, which includes a textured brain surface geometry, differentiates the corpus callosum, and includes the falx cerebri and tentorium cerebelli membranes. Because the EdiFEHM has not yet been validated in peer-reviewed literature, the key results from McGill are reproduced here.

##### Validation methodology

###### *Cadaver data.*

Three impact cases from Hardy *et al.*[4,5] were simulated. These cases—C755-T2, C288-T1, and C380-T2—span occipital, frontal, and temporal impact locations, with peak resultant linear accelerations ranging from 22g to 236g and peak rotational accelerations from 1,882 to 24,206 rad/s$^2$. The experimental kinematics were prescribed directly to the rigid skull of the FEHM. The simulated relative brain displacement was compared with displacement data recorded via neutral density targets (NDTs) embedded in the cadaver brain, and the simulated strain response was compared with strain histories recalculated from the NDT cluster displacement data by Zhou *et al.*[6]

###### *In vivo data.*

A low-severity rotational impulse from the tagged MRI experiments of Knutsen *et al.*[7] was also simulated. This dataset provides whole-brain strain distributions through axial, coronal, and sagittal planes, as well as the fractional volume of the brain exceeding strain thresholds of 0.02, 0.03, and 0.04.

###### *Assessment protocol.*

Quantitative comparison was performed using CORA (Correlation and Analysis, version 3.6.1),[8] with parameter settings optimised by Giordano and Kleiven,[9] for each validation criterion. These settings are reproduced in Supplementary Table 4. In addition to the CORA rating itself, an adapted biofidelity rating is proposed that averages the phase, magnitude and shape components of the CORA cross-correlation and scales the result to a 0–10 range, allowing classification according to bands adapted from those originally defined in ISO/TR-

9790 for anthropomorphic test devices. The biofidelity bands used are: Excellent ≥ 8.6, Good ≥ 6.5, Fair ≥ 4.4, Marginal ≥ 2.6, and Unacceptable < 2.6.[10,3]

**Supplementary Table 4 Optimised CORA (ver. 3.6.1) parameters for FEHM**

| Parameter | Intracranial Pressure | Rel. Brain Displacement | Brain Deformation |
|---|---|---|---|
| A_THRES | 0.030 | 0.030 | 0.030 |
| B_THRES | 0.075 | 0.075 | 0.075 |
| A_VAL | 0.010 | 0.010 | 0.010 |
| B_DELTA_END | 0.200 | 0.200 | 0.200 |
| T_MIN/MAX | 0.000/0.030 | 0.000/0.040 | 0.000/0.040 |
| D_MIN/MAX | 0.01/0.12 | 0.01/0.40 | 0.01/0.25 |
| INT_MIN | 0.70 | 0.80 | 0.70 |
| K_V/P/G | 3/1/1 | 3/1/1 | 3/1/1 |
| G_V/P/G | 0.33/0.33/0.33 | 0.33/0.33/0.33 | 0.33/0.33/0.33 |

Validation against cadaver data, as determined by Giordano & Kleiven.[9] Table reproduced from McGill.[3]

#### Results

##### *Relative brain displacement.*

The CORA ratings for the relative brain displacement response of the 'All' FEHM are summarised in Supplementary Table 5. The overall rating of 0.432 places the model on the boundary of the 'marginal' and 'fair' biofidelity classifications (adapted biofidelity scores of 5.63–5.72 across the brain geometry variants, corresponding to 'fair'). This is consistent with other FEHM in the literature, where relative brain displacement is known to be more sensitive to contact definitions and the representation of the cerebrospinal fluid (CSF) interface than the strain response.[3]

**Supplementary Table 5. CORA ratings for the relative brain displacement response of the EdiFEHM ('All' model)**

| Impact case | CORA rating |
|---|---|
| C755-T2 | 0.514 |
| C288-T1 | 0.447 |
| C380-T2 | 0.336 |
| **Overall** | **0.432** |

Compared with cadaver data from Hardy *et al.*[4,5]. The average rating across NDT displacement histories is reported for each impact, along with the overall rating across all three impacts. Reproduced from McGill.[3]

##### *Localised strain response.*

The CORA ratings for the localised brain strain response of the 'All' FEHM during the C380-T2 lateral impact are presented in Supplementary Table 6. The simulated 1st-principal and maximum shear strain histories within the element cluster surrounding NDT-4 in the right parietofrontal region were compared with strain values recalculated from empirical data by Zhou *et al.*[6] The overall CORA rating of 0.739 corresponds to a 'good' biofidelity classification, representing a substantial improvement over the 'marginal' displacement rating and indicating that the strain response is less sensitive to the contact and CSF simplifications present in the model.

**Supplementary Table 6. CORA ratings for the localised strain response of the EdiFEHM**

| Strain measure | CORA rating |
|---|---|
| 1st-principal | 0.774 |
| Maximum shear | 0.704 |
| **Overall** | **0.739** |

Ratings are for the EdiFEHM ('All' model) within the element cluster surrounding NDT-4 during the C380-T2 impact, compared with strains recalculated from cadaver data by Zhou *et al.*[6] Reproduced from McGill.[3]

##### *Spatial strain distributions.*

The spatial distribution of 1st-principal strain through axial, coronal, and sagittal brain slices was compared with tagged MRI data from Knutsen *et al.*[7] The 'All' FEHM produced strain distributions that most closely replicated the in vivo patterns among the model variants tested, including fragmented, asymmetric strain patterns characteristic of the empirical response. The peak 95th-percentile 1st-principal strain magnitude of 0.032 fell within the experimentally recorded range of 0.026–0.053. The fraction of the brain volume exceeding the highest strain threshold considered ($\varepsilon > 0.04$) also peaked within the empirical range. Full spatial distribution plots and volume fraction time histories are presented in McGill,[3] Chapters 6 and 7.

#### Summary

The EdiFEHM achieved a 'good' biofidelity rating (CORA = 0.739) for predicting localised brain strain against cadaver data and produced spatial strain distributions consistent with in vivo tagged MRI observations. These results indicate that the model provides a suitable representation of the human head for investigating brain strain responses to blunt impacts. Known limitations include the simplified tied-contact representation of the CSF-brain interface, which contributes to the lower displacement ratings, and the use of a single 50th-percentile male anatomy. For a comprehensive discussion of the model's development, validation, and limitations, the reader is referred to McGill.[3]
